# Contrasting Effects of Cytoskeleton Disruption on Plasma Membrane Receptor Dynamics: Insights from Single-Molecule Analyses

**DOI:** 10.1101/2024.09.09.612020

**Authors:** Leander Rohr, Luiselotte Rausch, Klaus Harter, Sven zur Oven-Krockhaus

## Abstract

Traditional models such as the fluid mosaic model or the lipid raft hypothesis have shaped our understanding of plasma membrane (PM) organization. However, recent discoveries have extended these paradigms by pointing to the existence of micro- and nanodomains. Here, we investigated the role of the cytoskeleton in general and whether the picket fence model, established in animal cells, is transferable to the plant cell system. By using single-particle tracking photoactivated localization microscopy (sptPALM) in combination with genetically encoded enzymatic tools for the targeted disruption of the cytoskeleton, we studied the dynamics and nanoscale organization of a selection of PM receptor-like kinases (RLKs) and receptor-like proteins (RLPs). Our findings show that the disintegration of actin filaments leads to decreased diffusion, more restrictive motion patterns, and enlarged clusters, whereas the disintegration of microtubules results in increased diffusion, more unconstrained diffusive behavior, and decreased cluster sizes of the tested RLKs and RLPs. These results underscore the potential unique regulatory functions of cytoskeleton components in plants and suggest an altered mechanism compared to the picket fence model of the animal cell system. Our qualitative data can serve as the foundation for further investigations aimed at developing a comprehensive and refined model of protein dynamics and organization in plant cells.

## Introduction

The plasma membrane (PM), together with the cell wall, functions as the first selective barrier between the cell and the environment (Gronnier et al., 2018; Jaillais and Ott, 2020). Singer and Nicolson (1972) emphasized the fundamental significance of biological membranes and their organization in general and proposed the widely known fluid mosaic model. Their assumption was that proteins can laterally diffuse within the membrane without major restrictions. However, this would result in uniformly distributed membrane-embedded proteins such as receptors, independent of the entire PM proteome, which is unequivocally not the case. One significant expansion of the model was the introduction of the “lipid raft” hypothesis, which suggests that lipid rafts, containing high levels of cholesterol and sphingolipids, serve as platforms with a high molecular order of proteins and lipids. These platforms facilitate, for instance, selective interactions between signaling proteins and effector molecules (Simons and Ikonen, 1997). The hypothesis found support in results from the mammalian field, which indicated a binary characteristic of membranes that are partitioned into detergent-resistant and detergent-sensitive fractions (Brown and Rose, 1992; Yu et al., 1973). Similar research was later conducted in plants, delivering comparable results (Borner et al., 2005; Laloi et al., 2007; Lefebvre et al., 2007; Mongrand et al., 2004; Morel et al., 2006). These studies showed that the detergent-resistant membrane protein profile is distinct from that of the whole PM. However, the isolation of detergent-resistant fractions and the detergents used may cause changes in the PM itself. Moreover, results derived from the application of newer methods, such as fluorescent microscopy techniques, raised the question of whether detergent-resistant membrane areas indeed define functional membrane rafts (Kusumi et al., 2005; Raffaele et al., 2009; Tanner et al., 2011). Although the existence of “lipid rafts” in plants, now referred to as “micro- or nanodomains”, is undoubtedly accepted, their dynamics, organization and regulation still require further research.

Besides the plant-specific family of remorin proteins serving as scaffolding factors (Jarsch and Ott, 2011; Raffaele et al., 2009), the asymmetric localization and order of lipids within the PM (Gronnier et al., 2018), as well as the cell wall-PM continuum (Martiniere et al., 2012), the cytoskeleton is believed to play a vital regulatory role in the organization of plant PMs (Jaillais and Ott, 2020). This hypothesis is based on the so-called picket fence model from the mammalian field (Kusumi et al., 2012). In this model, the cortical cytoskeleton, composed of actin filaments in mammalian cells, defines membrane domains by acting as a fence that restricts the lateral diffusion of lipids and proteins within these domains. The model postulates additional pickets, which are represented by transmembrane proteins that are anchored either by the cytoskeleton in the cytosol or the extracellular matrix. However, it is important to note that the model, as being based on animal cells, does not include specific properties of plants, such as the existence of cortical microtubules that could act as an additional fence (Jaillais and Ott, 2020). McKenna et al. (2019) demonstrated an impact of actin and microtubules on the diffusion of some but not all proteins in the PM. Moreover, additional studies revealed a change in or loss of nanodomain organization of proteins after cytoskeleton disruption, again for some but not for all tested proteins (Bücherl et al., 2017; Danek et al., 2020; Jarsch et al., 2014; Konrad et al., 2014; Lv et al., 2017; Raffaele et al., 2008; Szymanski et al., 2015). Chemicals such as latrunculin or oryzalin were used in these studies to disintegrate either the actin or the microtubule cytoskeleton. However, these chemicals are difficult to fine-tune in terms of their tissue-specific activity and concentration. In contrast, we utilized recently developed genetically encoded, enzymatic tools that mimic the function of those chemicals. Specifically, we employed the *Salmonella enterica* effector SpvB for actin cytoskeleton disintegration (Harterink et al., 2017; Vilches Barro et al., 2019) and a phosphatase-inactive variant of the atypical tubulin kinase PROPYZAMIDE-HYPERSENSITIVE 1 (PHS1ΔP) for microtubule cytoskeleton disintegration (Fujita et al., 2013; Vilches Barro et al., 2019).

To address the question of how the cytoskeleton may interfere with the dynamics and organization of plant PM proteins, we focused on a group of proteins associated with signaling mechanisms, namely the receptor-like kinases (RLKs) BRASSINOSTEROID INSENSITIVE 1 (BRI1), PHYTOSULFOKIN RECEPTOR 1 (PSKR1), FLAGELLIN-SENSITIVE 2 (FLS2) and BRI1-ASSOCIATED RECEPTOR KINASE (BAK1), along with the RECEPTOR-LIKE PROTEIN 44 (RLP44). RLP44 is proposed to be a cell wall integrity sensor that controls cell wall homeostasis through an interplay with BRI1 and its co-receptor BAK1. The three proteins form a ternary receptor complex in the PM of plant cells (Glöckner et al., 2022). Additionally, RLP44 has been associated with phytosulfokine signaling, as it forms a complex with the corresponding PSKR1 receptor and its co-receptor BAK1 (Garnelo Gomez et al., 2021; Holzwart et al., 2018). FLS2 was chosen as a protein that is not connected to RLP44. Although FLS2 interacts with BAK1, it has been demonstrated that this interaction takes place at a minimum distance of 11.1 nm from the BRI1-BAK1-RLP44 complexes (Glöckner et al., 2022).

We used SpvB and PHS1ΔP to analyze the cytoskeleton influence on the dynamic properties of the four above-mentioned PM receptors using single-particle tracking with photoactivated localization microscopy (sptPALM) in a transient *Nicotiana benthamiana* system. To gain insight into the underlying mechanisms, we focused our investigation on three key parameters: (i) The diffusion coefficients, (ii) the organization of the respective receptors into nanoscale-like protein clusters, and (iii) the classification of molecular movement into transient movement types. It is worth mentioning that the three parameters are not inherently linked, for example, a reduced diffusion coefficient does not necessarily result in larger nanoscale protein clusters and/or immobile movement patterns. Consequently, their separate evaluation can give insights into different regulatory mechanisms.

We were able to show a clear link between manipulated microtubule formation, reflected by an increase in the diffusion coefficient for most of the tested proteins. Conversely, disintegration of actin filaments predominantly led to reduced diffusion coefficients. We furthermore demonstrated that destroyed microtubules predominantly led to decreased cluster sizes of the analyzed proteins clusters, while actin destruction resulted in increased cluster sizes.

Additionally, the classification of the protein tracks into the motion types (i) free diffusion, (ii) confined diffusion, (iii) immobility and (iv) directed diffusion revealed another influence of the cytoskeleton. Our data show that proteins spend more time in free diffusion states in the absence of microtubules. Conversely, disintegration of actin filaments resulted in an overall more confined behavior.

The opposing effects on diffusion, cluster sizes and motion patterns demonstrate the potentially unique regulatory functions of cytoskeletal components in plants and suggest that the picket fence model is not directly transferable to plant systems. However, our research may provide a basis for further investigation to translate these findings into an extended or revised functional model.

## Results

### Genetically encoded, enzymatic tools for cytoskeleton disruption are functional

Initially, SpvB and PHS1ΔP (Vilches Barro et al., 2019) were modified for our needs to be applied in sptPALM experiments. To avoid potential compatibility problems with the used sptPALM fluorophores, we decided to generate versions of the genetically encoded enzymatic tools without fluorescent tags but instead with a hemagglutinin (HA) tag. In addition, the expression of SpvB and PHS1ΔP was designed to be under the control of constitutive promoters (Figure 1C and F). The modified protein tools were tested for their ability to destroy the integrity of the cytoskeleton by co-expressing them with corresponding marker proteins in *N. benthamiana*. To label actin filaments, we used actin-binding domain 2 (ABD2) of *Arabidopsis* fimbrin 1 fused to GFP at the C- and N- terminus (Wang et al., 2008). To test the state of the microtubules, we applied the MICROTUBULE-ASSOCIATED PROTEIN 65-8 (MAP65-8) fused to RFP, which is known to bind cortical microtubules (Smertenko et al., 2008). Epidermal cells of *N. benthamiana* leaves were investigated under the confocal microscope three days post infiltration (dpi) with the constructs-containing *Agrobacteria*. The clear actin filament and microtubule cytoskeleton disassembly was observed in the presence of the corresponding genetically encoded, enzymatic tool (Figure 1B and E), proving their applicability as disintegration tools in epidermal *N. benthamiana* leaf cells.

**Figure 1.**
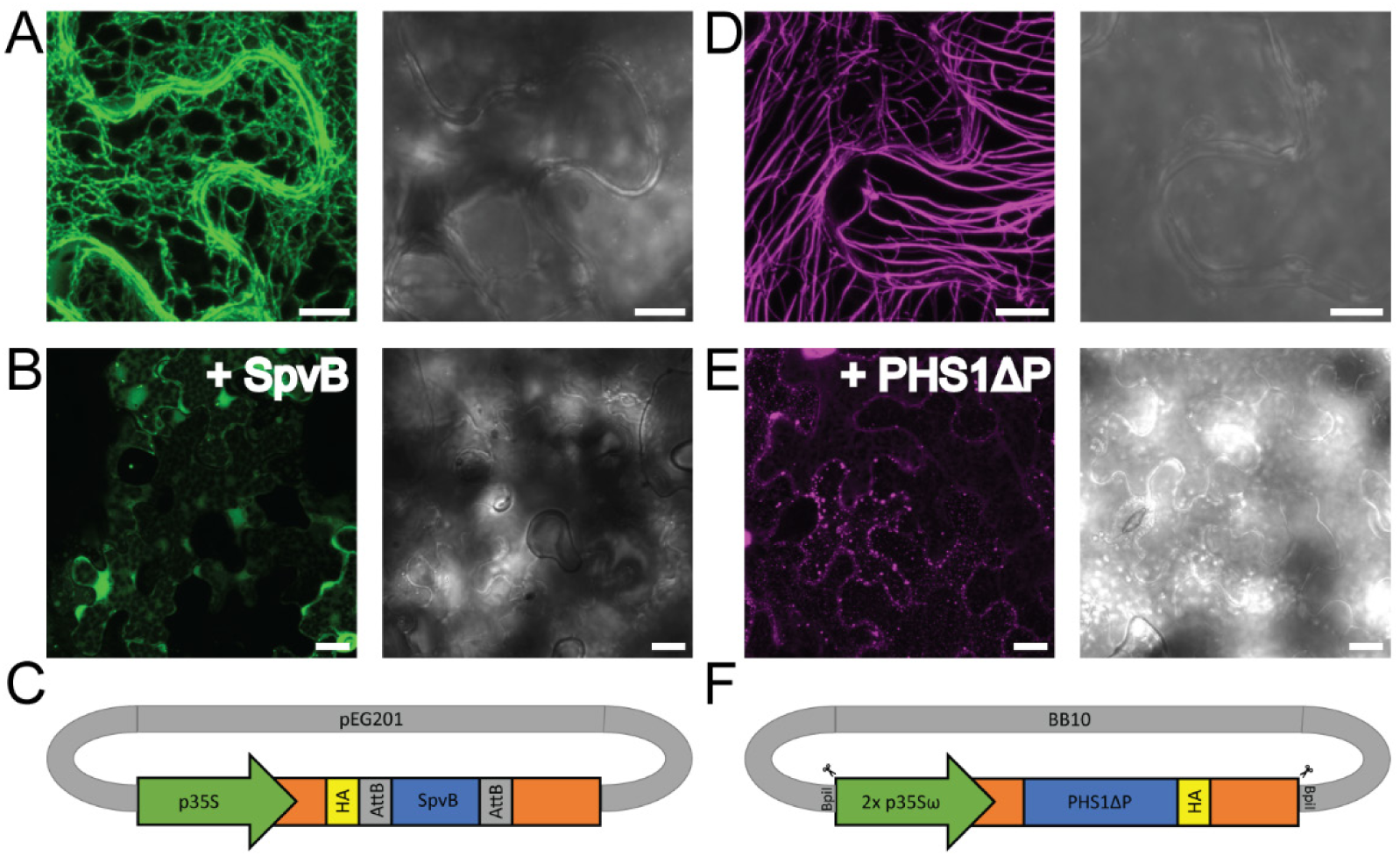
Overview of genetically encoded, enzymatic tools for cytoskeleton disintegration. **(A)** Exemplary confocal microscopy image of epidermal leaf cells of *Nicotiana benthamiana* (*N. benthamiana*) expressing the actin marker GFP-ABD2-GFP with the GFP channel on the left and the corresponding transmission light channel on the right. Intact actin filaments are clearly visible. Scale bar = 10 µm **(B)** Exemplary image of the co-expression of the disruption tool HA-SpvB and the actin marker GFP-ABD2-GFP in the GFP channel (left) and the corresponding transmission light channel (right). The co-expression with the disruption tool leads to removal of F-actin cables in all cells as shown before (Vilches Barro et al., 2019). Scale bar = 10 µm. **(C)** Schematic plasmid structure of the genetically encoded SpvB tool: By Gateway® technology SpvB was inserted into the pEG201 backbone (Earley et al., 2006) which contains a 35S promoter and an N-terminal HA-tag. **(D)** Exemplary confocal microscopy image of epidermal leaf cells of *N. benthamiana* expressing the microtubules marker MAP65-8-RFP with the RFP channel on the left and the corresponding transmission light channel on the right. Intact microtubules are observable. Scale bar = 10 µm. **(E)** Exemplary image of the co-expression of the disruption tool PHS1ΔP-HA and the microtubules marker MAP65-8-RFP in the RFP channel (left) and the corresponding transmission light channel (right). The co-expression with the disruption tool leads to the destabilization of cortical microtubules. Scale bar = 10 µm. **(F)** Schematic plasmid structure of the genetically encoded PHS1ΔP tool: The plasmid was generated by GoldenGate cloning (Binder et al., 2014) using Level I modules which were subsequently assembled in the Level II backbone of BB10. PHS1ΔP is under the control of a 2x 35Sω promoter module and fused C-terminally to an HA-tag. For the generation of higher order assemblies, BB10 contains Bpi I recognition sites.

### Disintegration of actin filaments predominantly leads to reduced protein mobility in the PM

To determine the influence of the actin cytoskeleton on the dynamics and the nanoscale organization of RLP44, BRI1, PSKR1, FLS2 and BAK1, we first expressed corresponding mEos3.2-tagged versions under the control of their native promoter in the absence or presence of HA-SpvB in *N. benthamiana* epidermal leaf cells. Subsequently, we used sptPALM and used the recently introduced OneFlowTraX software package for the analysis of protein dynamics and complex organization (Rohr et al., 2024). Independent of the co-expression with HA-SpvB, some general findings are worth mentioning: While the mEos3.2 fusions of RLP44, BRI1, PSKR1 and FLS2 showed one population of mobility each, BAK1-mEos3.2 presents a more confined and a more mobile variety (Figure 2A). The diffusion coefficients of RLP44-mEos3.2, BRI1-mEos3.2, PSKR1-mEos3.2 and FLS2-mEos3.2, as well as the one of the slower population of BAK1-mEos3.2, were comparable with other confined receptor proteins such as the PM intrinsic protein (PIP) 2;1 (D = 0,0047 µm²/s) reported before (Bayle et al., 2021; Hosy et al., 2015). In contrast, the more mobile variety of BAK1-mEos3.2 showed a diffusion coefficient about ten times higher than the other proteins (Figure 2B).

**Figure 2.**
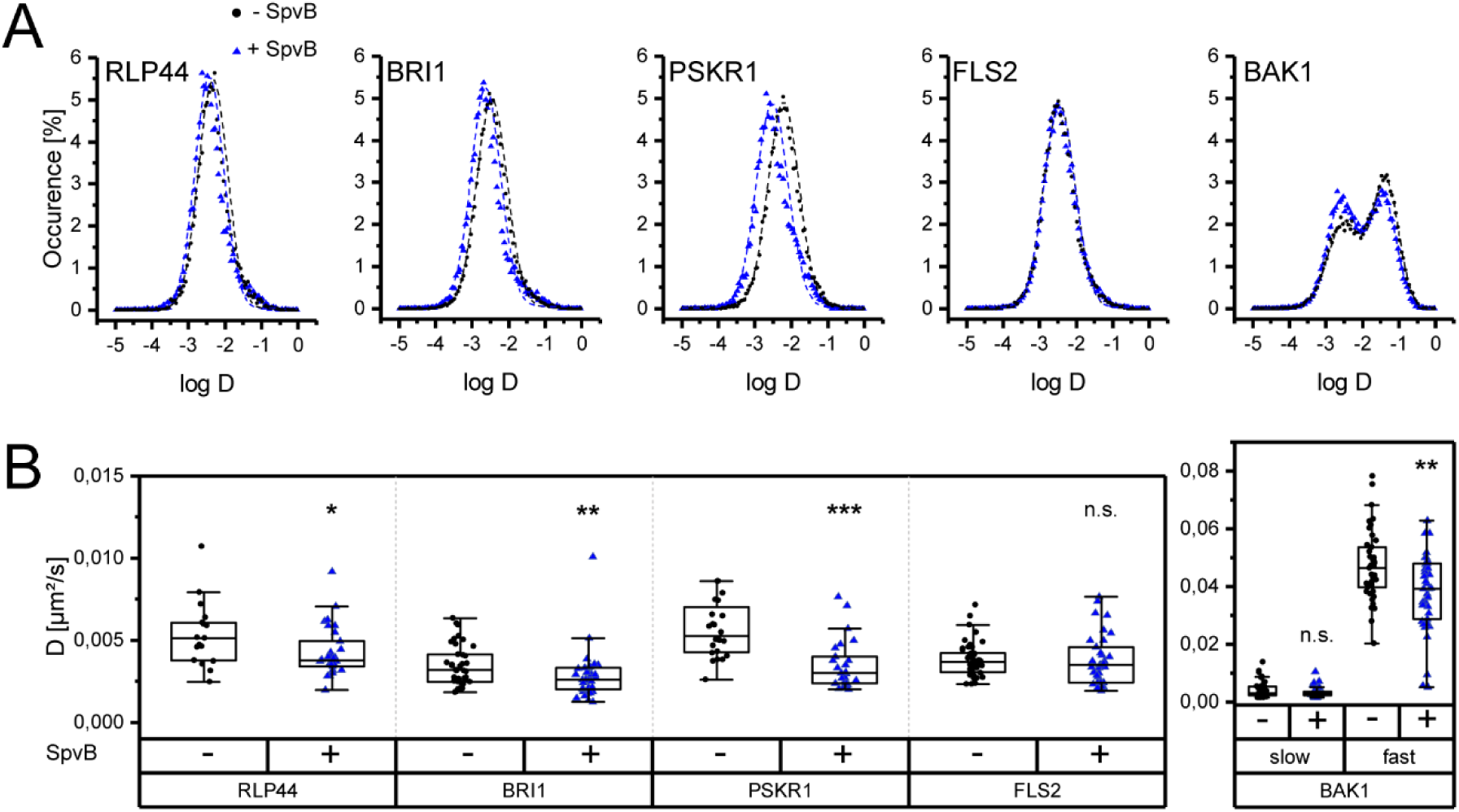
Disintegration of actin filaments primarily results in reduced protein dynamics in the plasma membrane. **(A)** Distribution of diffusion coefficients (D) represented as log(D) and plotted against their occurrence [%] over all quantified cells for RLP44-, BRI1-, PSKR1-, FLS2- and BAK1-mEos3.2. For each protein fusion, two distributions are shown: (i) In black, values obtained from epidermal *N. benthamiana* leaf cells expressing the respective fusion alone (- SpvB) and (ii) in blue values from the co-expression of the respective protein fusions with the genetically encoded, enzymatic tool SpvB (+ SpvB). For RLP44-, BRI1- and PSKR1-mEos3.2 a slight shift to lower log(D) values is observable, when SpvB is co-expressed. The effect is barely visible for FLS2-mEos3.2 and BAK1-mEos3.2. All measurements were performed three days post infiltration. Please note that all protein fusions show a bell-shape distribution (i.e., one mobility population), except for BAK1-mEos3.2 that presents a slower and a faster variety (two Gaussian fit). When co-expressed with SpvB, the slow fraction of BAK1-mEos3.2 is increased. **(B)** Representation of the peak D values of RLP44-, BRI1-, PSKR1-, FLS2- and BAK1-mEos3.2. with same color code as in (A). The peak values of individual cells (illustrated as single dots or triangles; n ≥ 17) were obtained by normal fits of distributions comparable to (A) expect of BAK1-mEos3.2 where a two-component Gaussian mixture model was applied (see Material *and Methods*). The separation of the BAK1 fractions was done according to this model with the peaks of the first maxima representing the slow fraction and the peaks of the second maxima the fast fraction, respectively. In the absence of intact actin filaments (blue; +SpvB) the diffusion coefficient is significantly decreased for the RLP44-, BRI1- and PSKR1-mEos3.2 fusions, while the reduction for FLS2-mEos3.2 is not significant. For BAK1-mEos3.2, only a decrease in the fast fraction is observable. For statistical evaluation, the data were checked for normal distribution and unequal variances and then analyzed according to the results of the test by applying either a Mann-Whitney U test or a One-way ANOVA. Whiskers show the data range excluding outliers, while the boxes represent the 25-75 percentile. p ≤ 0.001 (***); p ≤ 0.01 (**); p ≤ 0.05 (*); p > 0.05 (n.s.). All statistical analyses were performed with custom-made R scripts.

In the presence of HA-SpvB, we observed a significant reduction of the diffusion coefficient for RLP44-mEos3.2, PSKR1-mEos3.2, BRI1-mEos3.2 and the mobile population of BAK1-Eos3.2. In contrast, actin disruption did not affect FLS2-mEos3.2 dynamics. Moreover, the less mobile population of BAK1-mEos3.2 tended to have an even more reduced mobility. Additionally, the bimodal mobility distribution of BAK1-mEos3.2 shifted overall when the actin cytoskeleton was destroyed with HA-SpvB, increasing the fraction of the more restricted variant of BAK1-mEos3.2 compared to control cells.

### Disintegration of the microtubule cytoskeleton predominantly leads to enhanced protein mobility in the PM

While microtubules arise from centrosomes in animal cells, plants possess cortical microtubules (Farquharson, 2009). Their presence needs to be considered when a putative effect of the “membrane skeleton” on protein dynamics is investigated. Therefore, the identical set of RLK fusion proteins used for the actin approach (see above) were co-expressed with PHS1ΔP-HA, which specifically causes depolymerization of cortical microtubules (Fujita et al., 2013). While the mEos3.2-fusions of RLP44, BRI1, PSKR1 and FLS2 displayed one population for the diffusion coefficient, BAK1-mEos3.2 again showed two populations of different mobility, independent of the absence or presence of PHS1ΔP-HA (Figure 3A). In contrast to the data obtained after the destruction of actin filaments, the manipulation of cortical microtubules caused an increased diffusion of RLP44-mEos3.2 and PSKR1-mEos3.2. However, the mobility of BRI1-mEos3.2 and FLS2-mEos3.2 remained nearly unaffected in the presence of PHS1ΔP-HA (although a trend of increased diffusion was present for BRI1-mEos3.2). For BAK1-mEos3.2 a significant increase in the mobility was solely observed for the less mobile population (Figure 3B). Additionally, in the absence of polymerized microtubules, there was a slight shift in the BAK1 distribution between the two populations: More BAK1-mEos3.2 was present in the faster fraction and less in the slower one compared to the measurements where PHS1ΔP-HA was not present.

**Figure 3.**
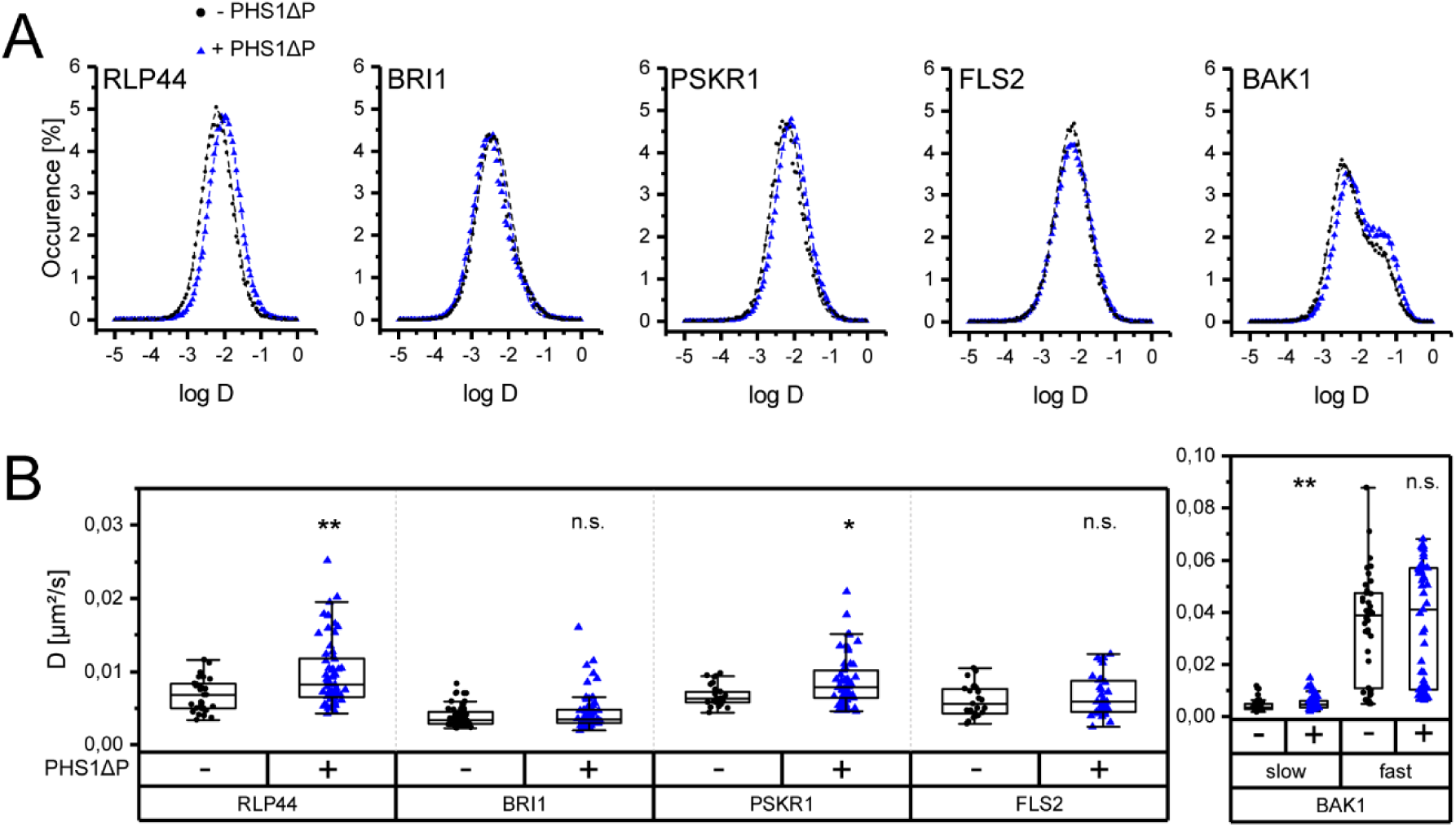
Disintegration of microtubule filaments primarily results in increased protein dynamics in the plasma membrane. **(A)** Distribution of diffusion coefficients (D) represented as log(D) and plotted against their occurrence [%] over all quantified cells for RLP44-, BRI1-, PSKR1-, FLS2- and BAK1-mEos3.2. For each protein fusion, two distributions are shown: (i) In black, values obtained from epidermal *N. benthamiana* leaf cells expressing the respective fusion alone (- PHS1ΔP) and (ii) in blue values from the co-expression of the respective protein fusions with the genetically encoded, enzymatic tool PHS1ΔP (+ PHS1ΔP). RLP44, PSKR1, and BAK1-mEos3.2 show a slight shift to higher log(D) values when co-expressed with PHS1ΔP. The effect is barely visible for the other protein fusions. Measurement conditions are as described in Figure 2. Again, all protein fusions show a bell-shape distribution (i.e., one mobility population), except for BAK1-mEos3.2 that presents a slower and a faster variety (two Gaussian fit). When co-expressed with PHS1ΔP, the fast fraction of BAK1-mEos3.2 is increased. **(B)** Representation of the peak D values of RLP44-, BRI1-, PSKR1-, FLS2- and BAK1-mEos3.2. with same color code as in (A) and obtained by n ≥ 25 cells. The evaluation was performed as described for Figure 2. In the absence of intact microtubules (blue; + PHS1ΔP) the diffusion coefficient is significantly increased for the RLP44- and PSKR1-mEos3.2 fusions as well as for the slow fraction of BAK1-mEos3.2, without a significant effect on the fast variety. Although BRI1- and FLS2-mEos3.2, as well as the fast variety of BAK1-mEos3.2, show no significant effect, a trend of increasing diffusion coefficients after microtubule disintegration is observable. Statistical analyses were conducted as in Figure 2, with the same box representation. p ≤ 0.001 (***); p ≤ 0.01 (**); p ≤ 0.05 (*); p > 0.05 (n.s.).

### The nanoscale organization of most PM proteins is changed by cytoskeleton disintegration

By manipulating the cytoskeleton structures, it is of great interest to know whether the nanoscale organization of protein clusters, commonly referred to as nanodomains, is affected as well (Jaillais and Ott, 2020; McKenna et al., 2019). To determine the cluster properties on the basis of spt data, several algorithms are available, such as Voronoi tessellation (Andronov et al., 2016; Levet et al., 2015), density-based spatial clustering of applications with noise (DBSCAN) (Ester et al., 1996) and the recently introduced nanoscale spatiotemporal indexing clustering (NASTIC) (Wallis et al., 2023). For our studies, we decided to apply the NASTIC algorithm since its approach is the least influenced by user-defined parameters and exclusively deals with track data as a whole (Rohr *et al*., 2024; Wallis et al., 2023). However, it is, like all the other algorithms, partly influenced by the set parameters and the raw data quality. Thus, we want to emphasize that only the relative changes in the nanoscale organization of clusters should be considered rather than absolute values.

Applying the NASTIC algorithm (see Material and Methods) to our tracking data in the absence or presence of actin-disintegrating SpvB, a predominantly decreased cluster diameter was observed for all tested PM proteins, with the exception of RLP44-mEos3.2 and FLS2-mEos3.2 (Figure 4A). In contrast, improper polymerization of cortical microtubules predominantly led to enlarged cluster diameters for all tested protein fusions, except for FLS2-mEos3.2, where the cluster size remained unaffected (Figure 4B).

**Figure 4.**
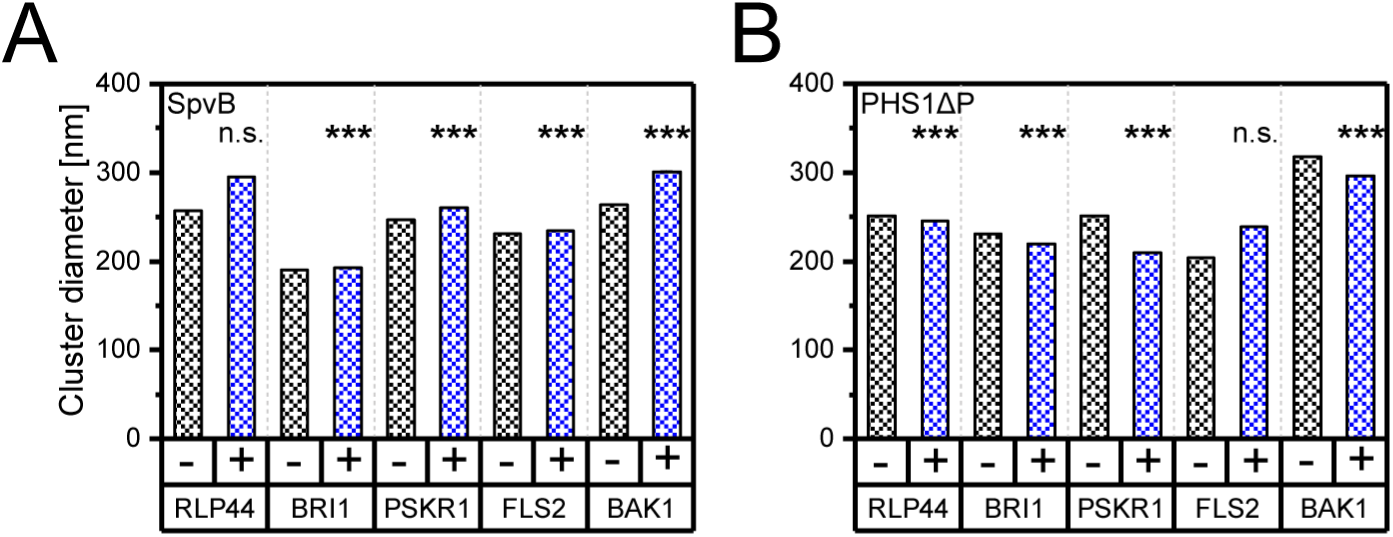
Actin and microtubules disintegration lead to opposing effects on nanocluster sizes. **(A)** Representation of corrected, transformed means of nanocluster diameter sizes (nm) of the tested protein fusions based on the track trajectories, also used for Figure 2 and Figure 3. The sizes were either determined for the protein fusions expressed alone (- SpvB, black) or together with the genetically encoded disruption tool for actin, SpvB (+ SpvB, blue). The analyses were performed by applying the NASTIC algorithm (Wallis et al., 2023) available in OneFlowTraX (Rohr et al., 2024) with a radius factor of 1.3 and at least three tracks per cluster. The corrected, transformed means are based on the log-normal distribution of all determined clusters among all evaluated cells (n as in Figure 2 and Figure 3), not considering clusters larger than 2,500 nm in diameter (≥ 1630). Except for RLP44-mEos3.2, all other tested fusions proteins show significantly increased cluster sizes upon the disintegration of actin filaments. Statistical analyses were performed according to Zhou et al. (1997). **(B)** Representation of corrected, transformed means of nanocluster diameter sizes (nm) as in (A) but here either with intact microtubules (i.e., expressed alone; - PHS1ΔP, black) or upon microtubules disintegration (+ PHS1ΔP, blue). Parameters and filtering as in (A) with ≥ 4356 evaluated cluster. Except for FLS2-mEos3.2, all other tested fusions proteins show significantly decreased cluster sizes upon the disintegration of microtubules filaments. Statistical analyses were performed as in (A). p ≤ 0.001 (***); p ≤ 0.01 (**); p ≤ 0.05 (*); p > 0.05 (n.s.).

In summary, the destruction of the actin filaments predominantly led to reduced mobility and smaller clusters, while the disintegration of the microtubule cytoskeleton resulted in higher mobility and enlarged clusters of most of the tested PM proteins.

### Classification of motion behavior reveals changes upon the disintegration of cytoskeleton components

In typical sptPALM experiments, the so-called short-range diffusion coefficient is extracted from molecular trajectories to obtain a value that is as independent as possible of directional motion, obstacles and boundaries (Saxton, 1997). Consequently, transient interactions of a diffusing protein with structural elements such as the cytoskeleton are not necessarily detectable by the diffusion coefficient alone. However, the long-term temporal evolution of these trajectories can be used to classify the type of movement (e.g., confined, directed, or free movement). For this purpose, several freely available software solutions and algorithms can be used (Das et al., 2009; Helmuth et al., 2007; Persson et al., 2013; Wagner et al., 2017).

Vega et al. (2018) introduced an efficient transient mobility analysis framework called “divide and conquer moment scaling spectrum” (DC-MSS), which was used to analyze the spatiotemporal organization of cell surface receptors and proved to be a suitable tool to analyze our data as well.

We focused on four major motion types of proteins, which are commonly used for studying biological systems. Proteins can (i) diffuse freely, for example in large unilamellar vesicles and in membrane blebs (Jaqaman and Grinstein, 2012; Kusumi et al., 2005), (ii) or become confined within structural corrals (Fujiwara et al., 2016). Furthermore, they can get (iii) anchored or immobilized when they bind to static components (Komura et al., 2016) or (iv) exhibit directed motion, for example, by transport processes via cytoskeleton components (Serge et al., 2003). Importantly, a protein may switch between different motion types during its lifespan. This was also suggested previously for plant PM proteins, such as PIP2,1 (Li et al., 2011).

In our analysis pipeline, the motion type classification was based on the same track data that were used before to evaluate the diffusion coefficients and cluster sizes. Individual tracks were classified with DC-MSS, and motion types were assigned either to entire tracks or track segments if the protein changed its motion type during the recording time. This allowed for an overall analysis of the relative time proteins spent in the following states: immobility, confined diffusion, free diffusion, and directed diffusion (Figure 5). A negligible number of tracks or track segments could not be classified (< 0.5 %) and were therefore excluded from the calculations.

**Figure 5.**
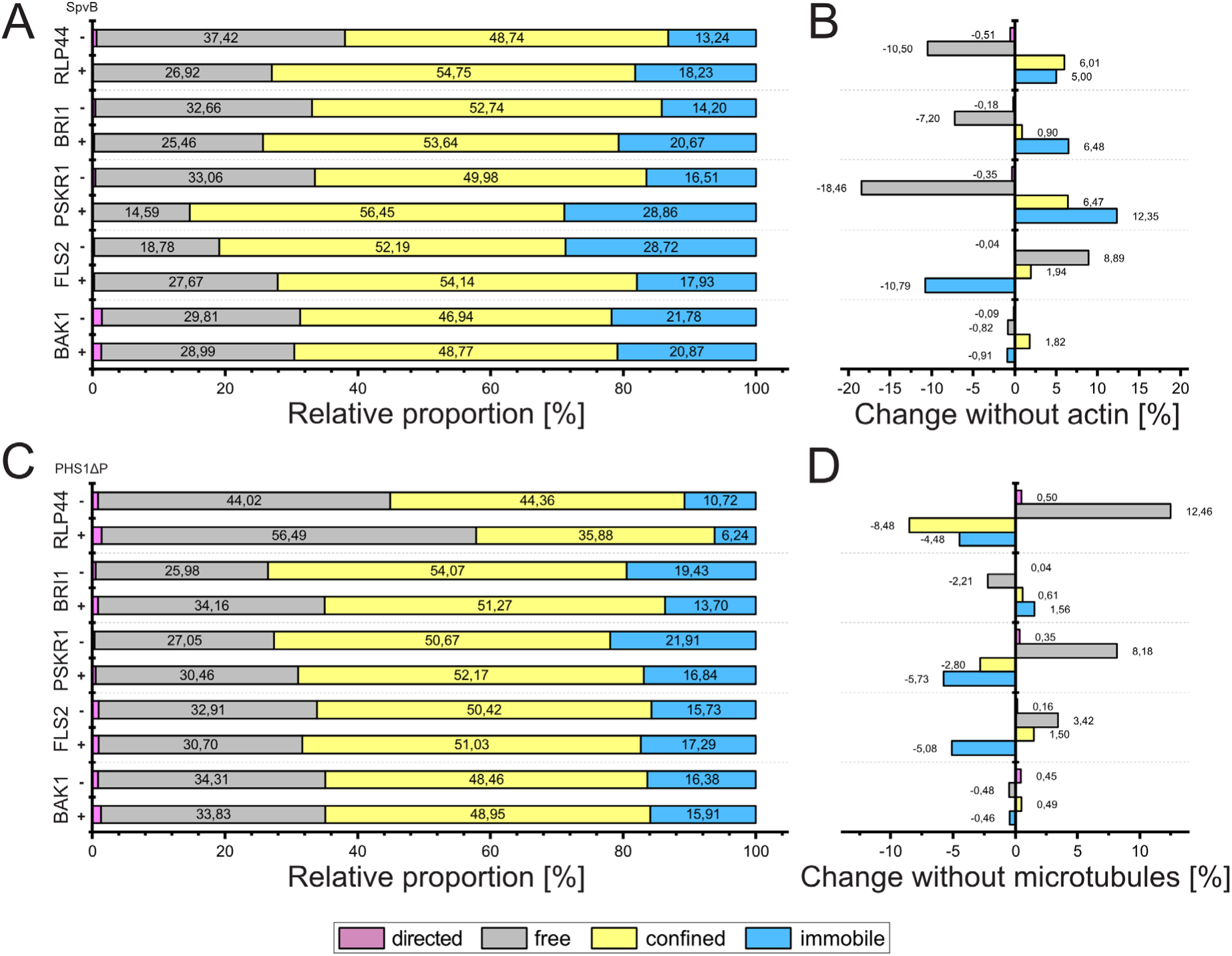
Classification of segment motion patterns depending on the cytoskeleton integrity. **(A)** Proportion of motion behavior (in percent) depending on the status of the actin cytoskeleton (either intact (- SpvB) or disintegrated (+ SpvB)), namely, directed (magenta), free (grey), confined (yellow) and immobile (blue). All studied proteins reveal primarily confined behavior, independent of the actin integrity. Immobility and free diffusive behavior are present in varying amounts amongst the proteins. Directed movement is barely present. The classification was performed according to Vega et al. (2018). **(B)** Relative shifts in the motion patterns in the absence of actin filaments with the same color code as in (A). RLP44-, BRI1- and PSKR1-mEos3.2 show a clear decrease in free diffusive behavior, while immobility and confined behavior (not for BRI1-mEos3.2) increases. However, FLS2-mEos3.2 shows a contrary effect by a decrease in immobility and an increase in free diffusive behavior. BAK1-mEos3.2 is nearly unaffected by the manipulation of the actin cytoskeleton. **(C)** Proportion of motion behavior (in percent) depending on the status of the microtubule cytoskeleton (either intact (- PHS1ΔP) or disintegrated (+ PHS1ΔP) with same behavior classes and color code as in (A) and (B). Again, all studied proteins reveal primarily confined behavior, independent of the microtubule disintegration except for RLP44-mEos3.2 (+ PHS1ΔP) that shows mostly free diffusive behavior. Immobility and free diffusive behavior are present again with varying amounts amongst the proteins, with distinct differences between RLP44-mEos3.2 (+ PHS1ΔP) and the other fusion proteins under manipulated conditions. Again, directed movement is barely present. **(D)** Relative shifts in the motion patterns in the absence of the microtubule cytoskeleton with the same color code as before. RLP44- and PSKR1-mEos3.2 show comparable effects with increased free diffusive behavior as well as with decreased confined movement and immobility. BRI1-mEos3.2 exhibits only minor changes while FLS2-mEos3.2 shows a slight increase in free and confined motion, while immobility is decreased, too. BAK1 is barely affected.

Under control conditions, RLP44-mEos3.2 mostly exhibited confined (49 %) and free behavior (37 %). In the presence of SpvB, i.e., in the absence of actin filaments, there was a decrease in the free diffusive behavior (- 11 %) while immobile and confined movement increased (+ 5 % and + 6 %, respectively). A comparable effect was observable for PSKR1-mEos3.2: Confined behavior represented the majority (50 %) under control conditions, and the disintegration of actin filaments by SpvB led to a shift in the motion patterns: The free diffusive motions decreased (- 18 %) while immobility and confined movement increased (+ 12 % and + 6 % respectively). These observations hold true for BRI1-mEos3.2 as well, although with less pronounced shifts as compared to RLP44 mEos3.2 and PSKR1-mEos3.2. In contrast, the consequences of actin disintegration on FLS2-mEos3.2 were contrary to those of the mEos3.2-fusions of BRI1, PSKR1 and RLP44. Here, a decrease in the immobile proportion (- 11 %) and an increase in the free diffusive behavior (+ 9 %) were detectable. Finally, BAK1-mEos3.2 exhibited a minor effect in response to actin filament disintegration, with shifts smaller than 2 %, suggesting a less eminent role of actin filaments on the motion behavior of BAK1 (Figure 5A and B).

As shown for the analysis of the diffusion coefficient and the cluster sizes, the disintegration of actin filaments and microtubules had contrary effects. This was also the case when evaluating the motion patterns.

In the presence of intact microtubules, RLP44-mEos3.2 mostly exhibited free diffusive or confined behavior (approx. 44 % each). However, with disintegrated microtubules, RLP44-mEos3.2 spent more time in a free diffusive state (+ 12 %) while confined and immobile behavior was less present (- 8 % and - 4 %, respectively). A comparable trend was observable for PSKR1-mEos3.2, where, under control conditions, confined motions represented the majority with 50 % followed by free diffusion (27 %). Again, with the co-expression of PHS1ΔP, there was in an increase in free diffusive behavior (+ 8 %), while immobile (- 5 %) and confined states (- 3 %) decreased. The relative changes for FLS2-mEos3.2 showed a decrease of immobility (- 5 %) in the absence of microtubules, a slight increase in the free diffusive proportion (+ 3 %) and a negligible effect on the confined behavior (+ 2 %). Compared to the strength of the shifts that RLP44-, PSKR1- and FLS2-mEos3.2 showed, BRI1’s changes were less pronounced, indicating a minor role of microtubules on the motion behavior of BRI1. This holds true as well for BAK1-mEos3.2, which exhibits shifts in the motion patterns below 1 % for all mobility classes (Figure 5C and D). All studied protein did not display substantial directed diffusive behavior, which suggests that the cytoskeleton components may not play a major role in transport processes (Figure 5).

In summary, we evaluated three parameters that gave insights into the organizational function of the cytoskeleton on PM proteins. Interestingly, diffusion coefficients, cluster sizes and motion classes were influenced in an opposite manner by the disintegration of actin filaments and microtubules. While the disintegration of actin filaments resulted predominantly in decreased diffusion but bigger clusters, the absence of microtubules increased the diffusion and led to smaller clusters. Concerning the motion classes, actin disruption predominantly caused a shift to more immobility and confined movement, while free diffusive behavior was reduced. Contrarily, the disintegration of microtubules resulted in increased free diffusive behavior, while immobility and the time proteins spent in confined states decreased. For BRI1 and BAK1, a less pronounced effect of the cytoskeleton on the motion behavior can be assumed, as the shifts between control conditions and the absence of actin filaments or microtubules were relatively low compared to the other PM proteins.

## Discussion

The PM plays a vital role for cell properties and a variety of biological processes. Its organization and associated functions have been the subject of several theories and considerations (Simons and Ikonen, 1997; Singer and Nicolson, 1972). Among them, the picket fence hypothesis is one of the most recent models that was initially studied in animal cells. Mainly due to the dynamic nature of the submembrane actin meshwork, no tractable experimental model for the mechanistic investigation of the fence or picket model is available. However, recently it was shown that actin rings in neurons compartmentalize the PM, acting as fences and confining membrane proteins (Rentsch et al., 2024). The direct transfer of the picket fence model from animal to plant cell biology is challenging since plant-specific properties need to be considered, such as the presence of cortical microtubules (Farquharson, 2009). Nevertheless, the model has also been tested and discussed as important for the dynamics and organization of plant PM protein dynamics and organization in several research articles and reviews (Jaillais and Ott, 2020; Martiniere et al., 2012; McKenna et al., 2019).

By applying genetically encoded enzymatic tools, we investigated the influence of the actin and microtubule cytoskeleton disintegration on the dynamics (i.e., via the diffusion coefficient and motion classification) and nanoscale organization of selected plant PM integral protein fusions, namely RLP44-mEos3.2, BRI1-mEos3.2, PSKR1-mEos3.2, FLS2-mEos3.2 and BAK1-mEos3.2, in the tobacco epidermal leaf cell system (Fujita et al., 2013; Harterink et al., 2017; Vilches Barro et al., 2019).

After the disintegration of the actin cytoskeleton by SpvB, we observed a decreased diffusion coefficient for all proteins, except for FLS2-mEos3.2. Furthermore, we exclusively detected two protein populations with different mobilities for BAK1-mEos3.2. The presence of such subpopulations usually indicates diverse molecular states of the protein. For instance, the subpopulation with lower mobility could be bound to cellular structures or other molecules, while the second one moves more freely. This may subsequently provide information about the biological roles of the protein (Hansen et al., 2018). Here, we speculate that the slow subpopulation of BAK1-mEos3.2 may be involved in signaling processes in restricted nanodomains, while the faster subpopulation is freely available for possible other interaction partners. Interestingly, the two subpopulations responded differently to actin filament disintegration via SpvB. Whereas no significant change was observed for the slower-moving BAK1-mEos3.2 population, the faster one showed a decreased diffusion comparable to the one of the other analyzed fusion proteins.

Furthermore, we observed that in the presence of intact actin filaments, the overall mobility distribution of BAK1 mEos3.2 features a higher fraction of the more mobile variant. The destruction of actin increases the relative amount of slower BAK1-mEos3.2, resulting in comparably equal amounts for both populations. Evidently, the functional meaning of these changes requires future experiments.

We additionally observed an increase in the cluster sizes for all tested proteins, except for RLP44-mEos3.2, after SpvB-mediated actin filament disintegration. This observation is in line with the picket fence model. In addition, the results can be integrated into a broader context: Recently, it was shown that the chemical destruction of actin leads to an increase in salicylic acid (SA) levels and the activation of SA-responsive genes in *A. thaliana* (Kalachova et al., 2019; Leontovycova et al., 2019; Matouskova et al., 2014). Moreover, the external application of SA constrained the diffusion of the PM auxin efflux carrier PIN-FORMED 2 (PIN2), followed by its condensation into PIN2 hyperclusters; a process mediated by remorins (Ke et al., 2021). We observed the same phenomenon for our fusion proteins: reduced diffusion with enlarged clusters. Whether these changes in the dynamics and nanoscale organization are related to SA requires further study.

However, so far, none of the available studies have shown a decrease in PM protein diffusion as a direct consequence of actin cytoskeleton disintegration (Hosy et al., 2015; McKenna et al., 2019).

For BRI1 and RLP44, no experiments have been published yet that tested the influence of the actin cytoskeleton on their dynamics. Interestingly, according to Lanza et al. (2012), brassinosteroid application modifies the actin cytoskeleton in an auxin-like manner by unbundling actin filaments. This implicates a contribution of BR signaling to the reorganization of the actin cytoskeleton. Moreover, a link between BR signaling and the actin cytoskeleton is further substantiated by the observation that the root-waving phenotype of the *Arabidopsis act2-5* mutant copies that of wildtype *Arabidopsis* seedlings treated with brassinolide (Lanza et al., 2012).

In contrast, the application of flg22, being the ligand of FLS2, increased the formation of actin filaments within three hours. This process is proposed to be the result of a well-orchestrated signaling cascade that (i) triggers the local high-order assembly of remorins that (ii) recruit formins, components comparable to mammalian integrins, which finally (iii) results in increased actin polymerization (Ma et al., 2021; Ma et al., 2022; Ma et al., 2023)

A long-standing paradigm that was still recently under debate was the direct involvement of actin during clathrin-mediated endocytosis. For example, it has been reported that FLS2 internalization depends on actin directly (Beck et al., 2012). However, Narasimhan et al. (2020) showed that actin is neither involved in membrane bending and scission nor in the initiation of endocytic processes. Rather, it is only required to transport endocytic vesicles after vesicle scission. Although this perspective seems to be acknowledged in the scientific community now (Kraus et al., 2024), current publications still refer to the old paradigm (Lu et al., 2023), stifling the debate. Whether the changed mobility and nanoscale organization of our tested proteins are a result of disturbed transport after vesicle scission due to actin filament disintegration requires further investigation.

In contrast to our results, McKenna et al. (2019) reported that the mobility of FLS2-GFP in epidermal hypocotyl cells of *A. thaliana* is enhanced after latrunculin treatment. However, we used SpvB, a bacterial effector from *Salmonella enterica,* as an actin disintegration tool. Although both latrunculin and SpvB are thought to specifically address actin, their modes of action differ: While SpvB acts almost exclusively on G actin by ADP-ribosylation (Hochmann et al., 2006), more recent publications indicate that latrunculin functions via direct binding to G actin (Spector et al., 1999) and can also affect F actin (Fujiwara et al., 2018). In addition, tool-specific, pleiotropic effects on cell physiology cannot be excluded. Both different modes of action and tool-specific site effects may have an impact on FLS2 dynamics. Another aspect that should be considered is that our experimental setup investigates proteins from *A. thaliana* in the heterologous cell environment of *N. benthamiana*. Therefore, the transfer of our results to cells of transgenic *A. thaliana* plants is not possible on a one-to-one basis, because the physiological, cellular and biochemical contexts of the *N. benthamiana* and *A. thaliana* cells are certainly not identical.

Besides studying the dynamics of the tested proteins via the diffusion coefficient, we analyzed their motion classes, too. We observed that all proteins show predominantly confined behavior under control conditions as well as in situations where the actin filaments were disintegrated. However, there was a substantial shift in the proportions of the motion classes for some but not all fusion proteins. In the absence of actin filaments, RLP44-mEos3.2 showed increased immobility and confined behavior, while free motion was reduced. The same holds true for PSKR1-mEos3.2 and BRI1-mEos3.2, although the effect on BRI1-mEos3.2 was less pronounced. In contrast, FLS2-mEos3.2 showed less immobile behavior but increased free motion in the absence of actin filaments. BAK1-mEos3.2 was nearly unaffected in its diffusive behavior when actin was disrupted.

In consideration of the combined results of the actin disintegration within the framework of the picket fence model, a direct transfer from the animal to the plant cell system does not appear to be possible without the inclusion of further aspects.

Due to the lack of actin filaments, an increase in the diffusion coefficient accompanied by enlarged nano-sized cluster structures and less confined and/or immobile behavior is expected (Fujiwara et al., 2002; Fujiwara et al., 2016; Jaqaman and Grinstein, 2012). However, our data prompt for a more sophisticated mechanism, as exclusively FLS2’s increase in cluster sizes and motion behavior in the absence of actin fits into the picket fence model. As actin is involved in a variety of processes, such as cell growth, cell division, cytokinesis, and various intracellular trafficking events (Szymanski and Staiger, 2018), pleiotropic effects cannot be excluded that are not linked to the role of actin in membrane organization. Additionally, in the absence of actin, a compensatory cytoskeletal interaction could take place through increased microtubule associations that would trap the proteins in confined corrals. Thus, a combined disruption of actin and microtubules will be of special interest in the future. So far, in the presented study, we focused on individual manipulations.

Through the disintegration of microtubules via PHS1ΔP (Fujita et al., 2013; Vilches Barro et al., 2019), we observed a contrary effect compared to that of the actin cytoskeleton disturbance: With the exception of BRI1-mEos3.2 and FLS2-mEos3.2, a significant increase in the diffusion coefficient was observed for the tested fusion proteins. Furthermore, reduced cluster sizes were observed for all tested receptors, again apart from FLS2-mEos3.2. Interestingly, for BAK1-mEos3.2, the significantly changed diffusion coefficient was this time observable for the slower subpopulation, while the faster one remained unaffected. This suggests a stronger connection between microtubules and the slower subpopulation of BAK1 than for the faster subpopulation.

Additionally, we analyzed the motion patterns of the fusion proteins. In the absence of microtubules, RLP44-mEos3.2, PSKR1-mEos3.2 and FLS2-mEos3.2 showed more free diffusive behavior, while BRI1- mEos3.2 and BAK1-mEos3.2 were barely affected.

In particular, the increased diffusion coefficients after microtubule destruction as well as the shift to more mobile motion patterns in combination with decreased immobility and confined motion are in good agreement with the picket fence model.

However, considering that the absence of physical barriers in the form of microtubules enables a less restricted motion of the fusion proteins, the decrease in cluster sizes after microtubule disintegration is difficult to interpret. Again, this change in nanoscale organization might be an indirect pleiotropic effect, as microtubules participate in a variety of processes in plant cells, such as the guidance of the cellulose synthase complexes to the PM (Paredez et al., 2006) or the maintenance of pavement cell morphogenesis (Belteton et al., 2018). Thus, alterations in the microtubule cytoskeleton organization may change the properties at the cell wall-PM interface that interfere with the dimension of the PM protein clusters.

Additionally, McKenna et al. (2019) showed that FLS2-GFP exhibits an enhanced diffusion coefficient after the disintegration of microtubules by oryzalin. In general, these observations are confirmed by our data, although in our case, the increase was less pronounced.

In summary, we studied three parameters that changed upon the disintegration of actin filaments and microtubules. Interestingly, the manipulated cytoskeleton components influenced the studied proteins in a contrary manner and resulted in distinct trends for diffusion coefficients, cluster sizes and motion patterns. While some aspects of either the actin or microtubule disruption match the picket fence model, others do not. Consequently, the model cannot be directly transferred from the animal field to the plant cell system. We conclude that this might be caused by the regulatory and functional responsibility of two cortical components, namely actin and microtubules, instead of cortical actin exclusively in animal cells. To further unravel a potential compensatory effect, the depletion of both structures will be a main task in the future. Additionally, studies in *A. thaliana* with inducible manipulations of the cytoskeleton will enable more background-free observation in the native organism.

We also started integrating our experimental data into a computational model generated by Smoldyn, a particle-based spatial simulation software (Andrews, 2009). Although modeling approaches have the potential to advance plant science to a great extent, the biggest challenge is still its underutilization in plant biology and thus the lack of comparative approaches (Dale et al., 2021). Recently, we predicted a new component of the fast brassinosteroid signaling pathway by computational modeling and emphasized its strengths in combination with wetlab experiments (Grosseholz et al., 2022). Based on our experience, we believe that the modeling of spatiotemporal dynamics may also reveal so-far hidden aspects in its regulation by the cytoskeleton.

## Material and Methods

### Plasmid construction

The genetically encoded enzymatic tools for the cytoskeleton manipulation, namely SpvB and PHS1ΔP, were provided by Prof. Alexis Maizel (COS Heidelberg) (Vilches Barro et al., 2019). For the generation of the expression constructs, the desired plasmid DNA was first amplified by PCR and then either a BP reaction into pDONR207 (for SpvB) or a blunt-end cloning reaction into pJET1.2 (Thermo Fisher Scientific) (for PHS1ΔP) was performed. Subsequently, an LR reaction was performed according to the manufacturer’s manual for SpvB into pEG201 (Earley et al., 2006) and cut-ligations with the needed Level I constructs into BB10 (Binder et al., 2014) were executed for PHS1ΔP. While SpvB cloning resulted in an N-terminally HA-tagged version under the control of the 35S promoter, PHS1ΔP is controlled by a 2x 35Sω promoter and C-terminally HA-tagged (Figure 1C and F). The BRI1-mEos3.2 and RLP44-mEos3.2 constructs are described in Rohr et al. (2024) and the construction of PSKR1-mEos3.2 was performed according to their protocol. BAK1-mEos3.2 and FLS2-mEos3.2 were provided by Dr. Birgit Kemmerling (ZMBP, Tübingen). The cytoskeleton markers GFP-ABD2-GFP and MAP65-8-RFP were kindly provided by Dr. Pantelis Livanos (FAU Erlangen).

### Plant material and growth conditions

All experiments conducted in this study were performed in transiently transformed *N. benthamiana* plants, cultivated under controlled greenhouse conditions. The desired proteins were transiently expressed using the AGL1 *Agrobacterium tumefaciens* strain (Lifeasible), as previously described (Hecker et al., 2015; Ladwig et al., 2015), without the washing step with sterile water. The plants were infiltrated with the respective constructs at an OD600 of 0.1 in a ratio of 1:1 or 1:1:1 with the silencing inhibitor p19. After watering, the plants were kept in ambient conditions and were imaged three days after infiltration.

### Confocal imaging

To confirm the functionality of the genetically encoded enzymatic tools for the cytoskeleton manipulation, the constructs were co-expressed with corresponding markers, namely GFP-ABD2-GFP (for actin) and MAP65-8-RFP (for microtubules). Subsequently, their localization inside epidermal leaf cells of *N. benthamiana* was investigated using confocal laser scanning microscopy on a SP8 laser scanning microscope (Leica Microsystems GmbH) with HyD detectors and a HC PL APOCS2 63 x/1.20 WATER objective three days post infiltration. For detection of the GFP signal, a 488 nm argon laser was used. The detection range was set to 500 nm – 550 nm. The images in Figure 1 are maximum projections that covered a z range of ∼ 15 µm obtained with a step size of 1 µm. The resulting images were processed with the help of the Leica Application Suite X (Version 3.3.0.16x). The detection of the RFP signal was performed as described above, using a detection range from 600 nm – 650 nm and a 561 nm diode pumped solid state laser.

### Sample preparation and movie acquisition for sptPALM measurements

All sptPALM measurements with transiently transformed *N. benthamiana* were performed three days post infiltration. For the acquisition, a small leaf area was cut out, excluding veins, and was placed between two coverslips (Epredia 24x50 mm #1 or equivalent) with a drop of water. The “coverslip sandwich” was then placed on the specimen stage, lightly weighted down by a brass ring to help flatten uneven cell layers. The composition of the custom-built microscope platform is described in detail in Rohr et al. (2024). For our purposes, the following filters for mEos3.2 were inserted into the emission beam path: 568 LP Edge Basic Longpass Filter, 584/40 ET Bandpass. The excitation power arriving at the sample was quantified (PM100D with S120C, Thorlabs) in epifluorescence mode after the objective to maintain consistency across experimental sets. Photoconversion of mEos3.2 was executed by applying moderate low intensities using 405 nm excitation. The signal of the red version of mEos3.2 was then obtained by excitation at 561 nm with 1800 µW. The magnification of the optical system was adjusted so that the length of one camera pixel corresponds to 100 nm in the sample plane. To identify feasible regions, larger areas of 51.2 x 51.2 µm were utilized by adjusting the focal plane and the VAEM angle with a frame rate of 10 Hz. In contrast, the recording was conducted in smaller regions of 12.8 x 12.8 µm and frame rates of 20 Hz by streaming between 2,500 and 5,000 frames per movie. A series of dark images were recorded at the same frame rate as the corresponding movies to correct for noise in data processing.

### Raw data processing and analysis of sptPALM movies

The experimental data sets were imported into OneFlowTraX (Rohr et al., 2024) to assess their quality. Samples exhibiting obvious outliers during the quality assessment were excluded from further analysis. The remaining files were analyzed using the “Batch Analysis” function of OneFlowTraX, according to the settings introduced by Rohr et al. (2024) for localization, tracking, and mobility analysis. The diffusion coefficients were calculated for all samples except BAK1 by fitting the data distribution with a Gaussian function and subsequently extracting the average diffusion coefficient from its peak center. For BAK1, two populations were clearly visible, necessitating the use of a two-component Gaussian mixture model to estimate their respective diffusion coefficients and relative fractions. The nanoscale organization of protein clusters was investigated using the NASTIC algorithm from Wallis et al. (2023) (also available in OneFlowTraX) with the following parameters: radius factor: 1.2 and at least three tracks per cluster. For the comparisons between the different cytoskeleton disintegration scenarios, the cluster diameter (nm) was used. This is calculated by treating the localizations in a cluster as a point cloud that is fitted by a two-dimensional Gaussian function. Its full width at half maximum (FWHM) values for x and y are then averaged to provide one value that is defined as the cluster’s diameter. The subsequent analysis steps were processed with custom-built R applications. Clusters with diameters greater than 2,500 nm were excluded from further analysis. Given that cluster data are log-normally distributed, specific statistical tests were employed to identify significant differences (Zhou et al., 1997). The respective figure (Figure 4) report the corrected, transformed mean as recommended by the aforementioned study.

### Motion classification with DC-MSS

The motion classification of individual (sub-)trajectories was conducted using the “divide-and-conquer moment scaling spectrum” (DC-MSS) algorithm (Vega et al., 2018). In short, trajectories that consist of at least 20 localizations are initially divided into segments of potentially disparate motion classes based on the extent of molecular movement. Subsequently, a movement scaling spectrum analysis is employed for the classification of these segments, utilizing threshold values to minimize the probability of misclassification among adjacent motion types. Intermediate refining steps are incorporated to enhance the confidence of both the trajectory segmentation and their classification.

## Acknowledgments and Funding

Our research was supported by the German Research Foundation (DFG) via the CRC 1101 (projects D02 and Z02) to S.z.O.-K and K.H. and by individual DFG grants to K.H. (HA 2146/22, HA 2146/23 We also thank the DFG for grants for scientific equipment (FUGG: INST 37/991-1, INST 37/992-1, INST 37/819-1, INST 37/965-1).

## Author contributions

Conceptualization, L.R. (Leander Rohr), K.H. and S.z.O.-K.; Data curation, L.R. and Ll.R. (Luiselotte Rausch); Formal Analysis, L.R. and Ll.R.; Funding acquisition, K.H.; Investigation, L.R. and Ll.R.; Methodology, S.z.O.-K.; Project administration, L.R., K.H. and S.z.O.-K.; Resources, Ll.R. and S.z.O.-K.; Software, S.z.O.-K.; Supervision, K.H.; Visualization, L.R.; Writing – original draft, L.R., K.H and S.z.O.-K.; Writing – review & editing, L.R., Ll.R., K.H and S.z.O.-K.

